# BART3D: Inferring transcriptional regulators associated with differential chromatin interactions from Hi-C data

**DOI:** 10.1101/2020.08.19.258095

**Authors:** Zhenjia Wang, Yifan Zhang, Chongzhi Zang

**Affiliations:** Center for Public Health Genomics, University of Virginia, Charlottesville, VA 22908, USA; Department of Biomedical Engineering, University of Virginia, Charlottesville, VA 22908, USA; Department of Public Health Sciences, and Department of Biochemistry and Molecular Biology, University of Virginia, Charlottesville, VA 22908, USA

## Abstract

**Summary:** Identification of functional transcriptional regulators associated with chromatin interactions is an important problem in studies of 3-dimensional genome organization and gene regulation. Direct inference of TR binding has been limited by the resolution of Hi-C data. Here, we present BART3D, a computational method for inferring TRs associated with genome-wide differential chromatin interactions by comparing Hi-C maps from two states, leveraging public ChIP-seq data for human and mouse. We demonstrate that BART3D can detect relevant TRs from dynamic Hi-C profiles with TR perturbation or cell differentiation. BART3D can be a useful tool in 3D genome data analysis and functional genomics research.

**Availability and Implementation:** Implemented in Python, source code freely available at https://github.com/zanglab/bart3d

**Supplementary Information:** Supplementary data are available.

## INTRODUCTION

The three-dimensional (3D) organization of eukaryotic genomes affects transcriptional gene regulation (Bonev and Cavalli, 2016; Gorkin *et al*., 2014; Yu and Ren, 2017). Although topologically associating domains (TADs) appear to be conserved across cell types at the level of cell populations (Rao *et al*., 2014; Schmitt *et al*., 2016; Dixon *et al*., 2012), chromatin architecture is highly dynamic during development, cell differentiation, or under experimental perturbation (Dixon *et al*., 2015; Li *et al*., 2015; Zheng and Xie, 2019), and can be disrupted in disease states (Bonev and Cavalli, 2016; Fang *et al*., 2020). Transcriptional regulators (TRs), including transcription factors and chromatin regulators, are required for the establishment and maintenance of chromosomal architecture (Steensel and Furlong, 2019; Stadhouders *et al*., 2018; Kim *et al*., 2016). Identification of functional TRs associated with chromatin dynamics can help unravel the spatial organization of the genome and the impact of 3D architecture on transcriptional regulation. The 3D organization of the genome can be measured using chromosome conformation capture-based methods such as Hi-C (Lieberman-Aiden *et al*., 2009) and in situ Hi-C (Rao *et al*., 2014). Chromatin interaction events can be detected by inferring loop structures from signal enrichment in Hi-C contact maps (Rao *et al*., 2014; Doyle *et al*., 2014). However, limited by the restriction enzyme digestion and ligation procedure and highly dependent on the sequencing depth, the resolution of Hi-C maps is typically 10^4^-10^5^ bp, or can be as high as 10^3^ bp for ultra-deep *in situ* Hi-C (Rao *et al*., 2014). It is still difficult to reach the sub-nucleosomal resolution of TR binding events (10^1^-10^2^ bp). HiChIP (Mumbach *et al*., 2016) and PLAC-seq (Fang *et al*., 2016) can reach higher resolution but require additional experimental steps to use a preselected protein factor as an anchor, limiting the feasibility for an unbiased TR association analysis. Computational models such as chi-CNN (Jaroszewicz and Ernst, 2019) can predict interactions at a high resolution, but require other data types such as DNase-seq and ChIP-seq as input in additional to Hi-C. It remains challenging to identify TR binding directly from low-resolution Hi-C data alone for functional analysis of 3D genome data.

Most computational methods for differential Hi-C data analysis, including diffHic (Lun and Smyth, 2015), FIND (Djekidel *et al*., 2018), HiCcompare (Stansfield *et al*., 2018), Selfish (Ardakany *et al*., 2019), and CHESS (Galan *et al*., 2020), focus on detecting changes in chromatin interaction events on the locus-to-locus level, i.e., differential loops. Few methods can generate a genome-wide differential interaction profile or make TR association analysis directly from differential Hi-C data. TR inference from collected genomic binding profiles is a more powerful approach than conventional DNA sequence motif search (Wang *et al*., 2018; Qin *et al*., 2020). We previously developed BART (Wang *et al*., 2018), an algorithm for inferring TRs whose binding profiles associate with a query genomic profile, leveraging over 13,000 human and mouse ChIP-seq datasets collected from the public domain. Here, we present BART3D, a new bioinformatics tool for 3D genome data analysis and TR inference integrating Hi-C maps with public ChIP-seq data.

## METHODS

The input of BART3D are Hi-C-type 3D genome contact maps from two biological states, with or without replicates for each state. BART3D first generates a genomic differential chromatin interaction (DCI) profile by comparing the contact maps, then uses the BART algorithm (Wang *et al*., 2018) to identify transcriptional regulators (TRs) whose binding sites are associated with either increased or decreased chromatin interactions (Fig. 1a). BART3D can accept three formats of unnormalized genomic contact maps as input: 1) raw count matrices from HiC-Pro (Servant *et al*., 2015), 2) .hic format files from Juicer (Durand *et al*., 2016), and 3) .cool format files (Abdennur and Mirny, 2020). The output includes ranked lists of TRs associated with increased or decreased chromatin interactions with a series of statistical measurements.

**Figure 1.**
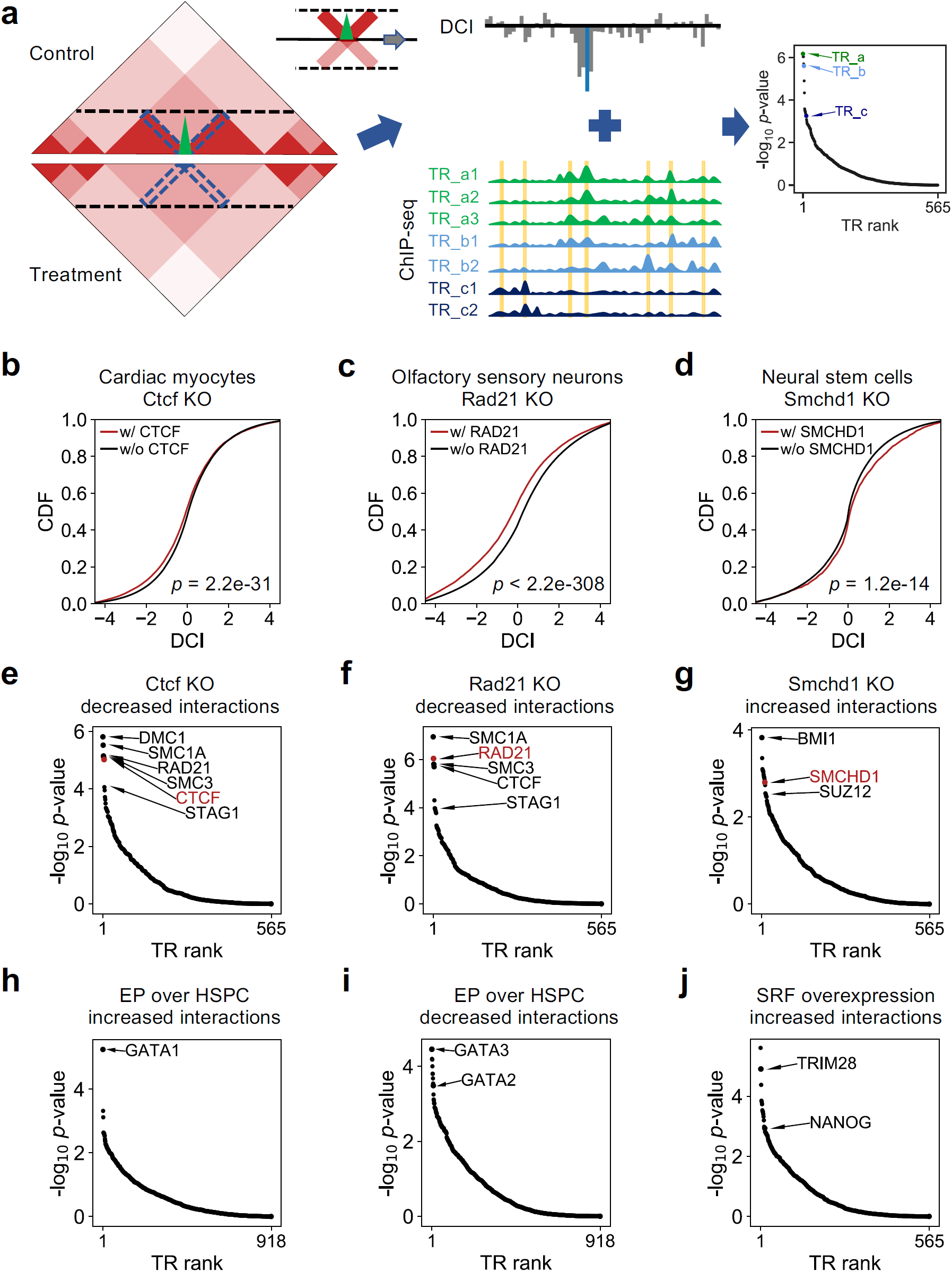
BART3D infers transcriptional regulators associated with differential chromatin interactions from Hi-C data. (**a**)BART3D workflow. BART3D takes Hi-C contact matrices from two biological states (Control and Treatment) as input, scans each chromosome to calculate the DCI score at every bin by comparing the interaction counts (blue dashed boxes at 45-degree angle) within a certain distance boundary (black horizontal dashed lines) between the two conditions, and derives a DCI profile. The BART algorithm is then applied to associate the DCI profile with a large collection of public transcription regulator (TR) ChIP-seq data for TR inference analysis. BART3D output is two ranked lists of all TRs associated with increase or decrease of chromatin interactions. (**b-d**) Cumulative distributions of DCI scores for genomic regions with (red) and without (black) binding sites of KO TRs, including Ctcf KO in cardiac myocytes (**b**), Rad21 KO in olfactory sensory neurons (**c**), and Smchd1 KO in neural stem cells (**d**), in mouse cell samples. DCI scores were calculated for each 5kb bin by comparing the normalized contact frequencies with its +/−200kb flanking regions. P-values were calculated by Wilcoxon rank-sum test. (**e-g**) BART3D results on decreased (**e, f**) or increased (**g**) chromatin interactions from corresponding Hi-C datasets in (**b-d**). P-value scores were calculated from rank sum using the null hypothesis under the Irwin-Hall distribution. Top ranked TRs were labeled, and the KO TRs were marked in red. (**h-j**) BART3D results on increased (**h**) and decreased (**i**) chromatin interactions comparing human erythroid progenitors (EP) with hematopoietic stem and progenitor cells (HSPC), and increased chromatin interactions after SRF overexpression in mouse neural progenitor cells (NPCs) (**j**).

We first employ an innovative approach to quantify the difference between Hi-C contact matrices from the two different conditions. To account for the negative correlation between intra-chromosomal interaction frequency and genomic distance (Supplementary Fig. S1) (Lieberman-Aiden *et al*., 2009) and to extract chromatin architecture information, we first normalize the contact matrix of each chromosome using a distance-based approach, where the read count in each bin pair is normalized by the average read count across all bin pairs at the same genomic distance (Supplementary Fig. S2). For any region in a chromosome, we consider the intra-chromosomal interactions between this region and its flanking regions within a certain genomic distance, e.g., 200 kb, quantified by a contact score array in the contact matrix, represented by the dashed, blue, 45-degree boxes in Fig. 1a. We calculate a DCI score for this region by comparing the two sets of contact score arrays from the two conditions, i.e., treatment and control. Specifically, we perform a paired t-test comparing each control contact score array with each treatment contact score array, and use Fisher’s method (Fisher, 1925) to calculate a combined statistic from all t-test p-values. The DCI score is then determined as logarithm of the p-value from the Fisher’s combined test, with the sign determined by the majority of the individual t-test statistics. Positive or negative DCI scores represent increased or decreased chromatin interactions, respectively, from control to treatment at this region (See Supplementary Methods for details). In this way, we generate a genome-wide DCI profile by scanning all chromosomes to calculate DCI scores for all non-overlapping bins across each chromosome.

We then infer TRs whose genome-wide binding profiles are associated with the DCI profile derived from Hi-C contact matrices. We map the genomic DCI profile to the union DNaseI hypersensitive sites (UDHS), a curated dataset representing all putative cis-regulatory elements (CREs) in the genome (Wang *et al*., 2016), and generate a cis-regulatory profile in which the score for each candidate CRE is set to equal the DCI score of the genomic region in which the CRE is located. We use the BART algorithm (Wang *et al*., 2018) to infer TRs that preferentially bind at CREs with a high score, representing increased chromatin interactions. Then we flip the cis-regulatory profile and perform BART analysis again, to infer TRs whose binding are associated with decreased chromatin interactions (Fig. 1a).

## RESULTS

To demonstrate the performance of BART3D, we calculated DCI profiles and inferred TRs for several published Hi-C experiments comparing wild type (WT) with DNA-associating factor knockout (KO) models in mouse cells. The KO targets include transcription factor Ctcf (Rosa-Garrido *et al*., 2017) and cohesin complex component Rad21(Canzio *et al*., 2019), which are known to function cooperatively to induce DNA looping and maintain TAD structures (Fudenberg *et al*., 2016), as well as Smchd1 (Jansz, Keniry, *et al*., 2018), which has repressive effects on transcriptional regulation and chromatin architecture. As expected, genomic regions containing binding sites of CTCF or RAD21 exhibit decreased chromatin interaction levels after KO (Fig. 1b,c), while those containing SMCHD1 sites associate with increased chromatin interactions after KO in their corresponding samples (Fig. 1d). This result indicates that the DCI profile can connect perturbed protein binding sites with differential chromatin interaction. Indeed, the BART3D results show that the KO factors are always among the top ranked TRs inferred to be associated with the corresponding decreased or increased chromatin interactions (Fig. 1e-g, labeled in red). These results show that BART3D can successfully infer TRs that induce chromatin interaction changes from Hi-C data.

In addition to the KO factor itself, we also found other TRs highly ranked in the BART3D results from the KO/WT Hi-C comparisons (Fig. 1e-g). For Ctcf or Rad21 KO (Fig. 1e,f), several top inferred TRs, including SMC1A, SMC3, and STAG1 are all components of the cohesin complex (Peters *et al*., 2008). For Smchd1 KO (Fig. 1g), BMI1 and SUZ12 are related to polycomb group (PcG) factors, which have been shown to interact with SMCHD1 (Jansz, Nesterova, *et al*., 2018) and have repressive effects on transcription and chromatin state.

Besides TR KO data, we further tested the ability of BART3D to infer functional TRs from Hi-C data across cell types during differentiation. Comparing Hi-C data between human hematopoietic stem and progenitor cells (HSPCs) and differentiated erythroid progenitors (EPs) (Zhang *et al*., 2020), GATA1, which is specifically expressed in erythroid development (Leonard *et al*., 1993), ranked on the top of BART3D result on EP increased chromatin interactions (Fig. 1h). The top ranked TRs associated with EP decreased (high in HSPC) chromatin interactions (Fig. 1i) include GATA3, a transcription factor required in the maintenance of hematopoietic stem cells (Ku *et al*., 2012), and a key HSPC-associated factor GATA2 (Zhang *et al*., 2020). We also compared Hi-C data in mouse neural progenitor cells (NPCs) before and after overexpression of Srf, a transcription factor repressing cell-type-specific genes and promoting cellular reprogramming to pluripotency (Ikeda *et al*., 2018). The top ranked factors associated with Srf overexpression-increased chromatin interactions reported by BART3D include TRIM28 and NANOG, both of which are involved with maintaining pluripotency of stem cells (Seki *et al*., 2010) (Fig. 1j). These results indicate that BART3D can identify relevant TRs from differential Hi-C analysis and can provide functional insights into the relations between chromatin architecture dynamics and transcriptional regulation during cell differentiation.

## DISCUSSION

We developed BART3D for differential analysis of Hi-C data and to infer functional TRs associated with changes in chromatin interactions. BART3D overcomes the relatively low resolution of Hi-C data and connects chromatin interactions on the multi-kb to Mb level to cis-regulatory events on the nucleosomal or base-pair level by accounting for statistical differences in Hi-C signals within a large distance range and using a predefined genomic CRE set. BART3D uses a distance-based normalization approach, which can remove cross-sample biases (Supplementary Fig. S2) and outperforms ICE normalization (Servant *et al*., 2015) in detecting local chromatin interactions (Supplementary Fig. S3). We use dynamic Hi-C datasets from TR KO experiments and cell differentiation to show that BART3D can infer TRs inducing chromatin architecture changes and other TRs with biological relevance.

In the framework of BART3D, we assume that differential chromatin interactions mainly associate with genomic binding of transcriptional regulator proteins, which act primarily in cis. Other events that can also result in pattern changes on Hi-C maps such as genome rearrangements are not considered in BART3D. Under this assumption, we focus on intra-chromosomal interactions within a certain range of chromosomal distance and ignore inter-chromosomal interactions. The default genomic distance is set as 200 kb, but users can adjust this parameter in exploratory studies for optimizing discovery power, as different TRs may associate with chromatin interactions at different genomic ranges (Supplementary Fig. S4). There are not many other tunable parameters. The bin size should be consistent with the Hi-C contact maps under interrogation and is restricted to the Hi-C data resolution. While replicates of Hi-C data have been accounted for in the DCI calculation, they can also be used to generate a background control for TR inference, i.e., TRs inferred from comparing replicates of Hi-C data from the same biological condition are likely due to technical variations and can be considered as false positive. Such TRs should be discarded if they also appear in results from cross-condition Hi-C comparisons.

Although developed for Hi-C data analysis, BART3D can also be applied to other 3D-genome data, such as ChIA-PET (Fullwood *et al*., 2009), HiChIP (Mumbach *et al*., 2016), and PLAC-seq (Fang *et al*., 2016). When analyzing HiChIP or PLAC-seq data using BART3D, one may notice that the ChIP factor tends to appear on the top of the inferred TR list. Because HiChIP/PLAC-seq signals are always enriched at genomic binding sites of the ChIP factor regardless of chromatin interaction changes, the inference of a ChIP factor and its known co-factors should be considered false positives and removed for result interpretation. We plan to account for this effect and develop an extended version for analyzing HiChIP/PLAC-seq data in the future. Because of the model design, BART3D is limited in detecting TRs that only associate with individual or a very small number of interacting loci in the genome. The normalization procedure in BART3D only accounts for the genomic distance, and should not replace other normalization approaches when pre-processing the data. For example, potential biases in Hi-C data such as GC content, sequence mappability, chromatin accessibility, and restriction enzyme cleavages can be considered using other normalization methods (Yaffe and Tanay, 2011; Hu *et al*., 2012). In addition, TR inference in BART3D is limited to collected ChIP-seq data, which currently include 918 human TRs and 565 mouse TRs but still grow rapidly and require regular updates and maintenance. Nevertheless, BART3D provides a framework for accurate inference of TRs associated with differential chromatin interactions and has broad applications in making biologically meaningful inferences and generating hypotheses from 3D genome data.

## FUNDING

This work was supported by the US National Institutes of Health [R35GM133712, K22CA204439] and a Phi Beta Psi Sorority Research Grant to C.Z.

## ACKNOWLEDGMENTS

The authors would like to thank Dr. Panagiotis Ntziachristos for helpful discussions and members of the Zang Lab for critical reading of the manuscript and testing the software.

## Conflict of Interest

none declared.

## Notes

https://github.com/zanglab/bart3d

### Competing Interest Statement

The authors have declared no competing interest.

