## Supplementary data for "BART3D: Inferring transcriptional regulators associated with differential chromatin interactions from Hi-C data"

Supplementary data include Supplementary methods, 4 Supplementary figures, and 3 Supplementary tables.

**Supplementary Figure S1.** Hi-C read count negatively correlates with genomic distance between bin pairs.

**Supplementary Figure S2.** Effect of normalization.

**Supplementary Figure S3.** Comparison of different normalization methods.

**Supplementary Figure S4.** BART3D results on dynamic Hi-C datasets with TR perturbation under different genomic distance settings.

**Supplementary Table S1.** List of Hi-C data used in this work.

**Supplementary Table S2.** List of ChIP-seq data used in this work.

**Supplementary Table S3.** List of HiChIP data used in this work.

### SUPPLEMENTARY METHODS

#### Data collection

Hi-C, HiChIP, and ChIP-seq data were collected from NCBI GEO (Barrett *et al.*, 2013) in fastq format. Detailed information including accession numbers of all samples used in this work can be found in Supplementary Tables S1-S3.

#### Data processing

Hi-C (Supplementary Table S1) and HiChIP (Supplementary Table S3) sequence reads were aligned to the human (hg38) or mouse (mm10) reference genomes and processed using HiC-Pro (Servant *et al.*, 2015). Contact matrices were generated at a resolution of 5kb and normalized as described in *Normalization of contact matrices*. ChIP-seq (Supplementary Table S2) reads were aligned to the mouse reference genome (mm10) using BWA (Li and Durbin, 2009). Sam files were then converted into bam files using samtools (Li *et al.*, 2009). MACS2 (Zhang *et al.*, 2008) was used to call peaks under the FDR threshold of 0.05.

#### Normalization of contact matrices

Given a Hi-C contact matrix $A = \{a_{ij}\}$, the observed read count $a_{ij}$ represents the interaction frequency between a pair of genomic bins $i$ and $j$. To account for the negative correlation between the intra-chromosomal interaction frequency and the genomic distance between the bin pair (Lieberman-Aiden *et al.*, 2009), we normalized the contact matrix of each chromosome as follows: for any given genomic distance $d_k = k * r$, where $r$ is the bin size (data resolution), we employed a normalization factor $\bar{S}_{d_k}$ as the average read count across all bin pairs with the same genomic distance $d_k$ in this chromosome, i.e., $\bar{S}_{d_k} = (\sum_{j-i=k} a_{ij})/n$, where *n* is the total number of bin pairs with distance $d_k$. The read count $a_{ij}$ of the bin pair with distance $d_k$ was normalized by $\bar{S}_{d_k}$ as $a'_{ij} = a_{ij}/\bar{S}_{d_k}$. The matrix $A$ was normalized into $A' = \{a'_{ij}\}$ for each chromosome.

**Detection of differential chromatin interactions**

Considering *m* Hi-C matrices for treatment and *n* Hi-C matrices for control (m,n≥1) as input, we denoted the normalized matrix $T^i = \{t^i\}$ as the *i*-th treatment matrix and $C^j = \{c^j\}$ as the *j*-th control matrix, (*i*=1, …, *m*; *j*=1, …, *n*). $\mathcal{B} = \{1, 2, \ldots, \lfloor l/r \rfloor\}$ represents all equal-sized non-overlapping bins within a chromosome, where $l$ is the length of the chromosome and $r$ is the bin size. For a given genomic region $x \in \mathcal{B}$ and a pre-defined range of genomic distance $L$, the interaction frequencies between $x$ and its flanking regions with genomic distance up to $L$ were collected, as $IT^i = \{t^i_{xk}\}$ from $T$ and $IC^j = \{c^j_{xk}\}$ from $C$, respectively, where $k \in \mathcal{B}, x - L/r \leq k \leq x + L/r$. The t-statistic at $x$ was calculated using the paired-sample *t*-test between the two arrays of interaction frequencies $IT^i$ and $IC^j$ as follows:

$$d_{xk} = t^i_{xk} - c^j_{xk},$$

$$t^{ij} = \frac{\bar{d}}{s_d/\sqrt{n}}$$

where $d_{xk}$ is the difference in interactions between each paired element in $IT^i$ and $IC^j$; $\bar{d}$ and $s_d$ are the mean and standard deviation of $\{d_{xk}\}$, respectively; $n$ is the length of each array and $n = 2L/r + 1$.

The estimated p-value of $t^{ij}$ is $p^{ij}$.

Assume we have $k^+$ p-values estimated from positive t-statistics and $k^-$ p-values estimated from negative t-statistics, $k^+ + k^- = mn$. We use Fisher's method (Fisher, 1925) to combine all the $k^+$ p-values as:

$$\chi^2 = -2\sum_{i=1}^{k^+} \ln(p^i)$$

The statistic $\chi^2$ follows a chi-squared distribution with $2k^+$ degrees of freedom. Under this statistical distribution, a p-value can be determined as $p^+$ to quantify the significance of chromatin interaction increases between multiple treatment and control matrices at the genomic region x. Meanwhile, the $k^-$ p-values estimated from negative t-statistic can be combined as $p^-$ using the same approach. The differential chromatin interaction (DCI) score at the given genomic region can be calculated using the logarithm of $p^+$ or $p^-$ as follows:

$$DCI = \begin{cases} -\log_{10}(p^+), if\ k^+ > k^-, or\ k^+ = k^-;\ p^+ < p^- \\ \log_{10}(p^-), if\ k^+ < k^-, or\ k^+ = k^-;\ p^+ > p^- \end{cases}$$

**Inference of TRs associated with differential chromatin interactions**

We used previously curated union DNaseI hypersensitive sites (UDHS), which include 2,723,010 unique non-overlapping DNase-seq peaks for human and 1,529,448 for mouse, to represent all putative cis-regulatory elements (CREs) in the genome (Wang *et al.*, 2016). A genome-wide DCI profile was generated by calculating the DCI score of every bin across each chromosome. The DCI profile was mapped to UDHS such that the score for each candidate CRE is set to be equal to the DCI score of the genomic bin where the CRE is located.

We used the BART algorithm (Wang *et al.*, 2018) to infer TRs associated with differential chromatin interactions. The analysis was done twice, for inferring TRs associated with increased and decreased chromatin interactions, separately. For increased chromatin interactions, we ranked all CREs decreasingly by their scores, i.e., CREs with high positive scores would be ranked at the front. We calculated an association score between the CRE profile and each TR binding profile for all ChIP-seq datasets. The association score is defined as the area under the ROC curve (AUC) using the DCI score on CRE as the predictor for TR binding, set as a binary value indicating whether the CRE is overlapped with a peak of that TR from the ChIP-seq dataset. To account for multiple ChIP-seq datasets for the same TR, the Wilcoxon rank-sum test was then applied to assess each TR's significance by comparing the association scores from all ChIP-seq data for this TR with those from all other ChIP-seq datasets, and a background model was used to detect the specificity of each TR. A series of quantification scores with statistical assessments were included for a final ranked list of inferred TRs. For decreased chromatin interactions, the CRE profile was flipped, so that the CREs with the most decreased chromatin interactions are ranked at the front, and the BART analysis was then performed in the same way.

**Comparison of normalization methods in detecting chromatin interactions**

To evaluate the feasibility and performance of HiC normalization approaches, we used HiChIP data and tested how different normalization methods affect the inference of the HiChIP target factor. By targeting a specific factor of interest, HiChIP signals are enriched at the target-bound loci (Mumbach *et al.*, 2016). We collected 84 HiChIP datasets targeting different TRs (Supplementary Table S3). For each HiChIP dataset, we generated genomic contact maps at 5kb resolution without normalization, with ICE normalization (Servant *et al.*, 2015), and with distance-based normalization. We then generated a genomic profile from each contact map, in which each 5kb bin across the genome is scored as the sum of interaction signals between this bin and all of its flanking bins within 500kb. We used BART to infer TRs associated with this genomic interaction profile. We expected that a contact map with appropriate normalization should yield to a BART result in which the HiChIP target factor ranked higher (with higher significance). As a control, we run BART analysis on the HiChIP sequence read pile-up profile and used the rank of the target factor as a reference (Supplementary Fig. 3).

**Determination of default genomic distance parameter**

Using different genomic distance range parameters might lead to different TR inference results, because the acting range of different TRs vary a lot. In practice, users may try different distance parameters for exploratory studies. To set an appropriate default value for this parameter, we applied BART3D on a series of Hi-C datasets (Supplementary Table S1) comparing the wide type with perturbation (deletion or activation) of different TRs using different genomic distance ranges, i.e., 50kb, 100kb, 200kb, 500kb, and 1000kb. We compared the rank of the perturbation target factor in the BART3D results across different genomic distance ranges, and found that 200kb is where most perturbation factors were ranked on top (Supplementary Fig. S4). Therefore, we set 200 kb as the default value for the genomic distance range parameter.

**Supplementary Figure S1**

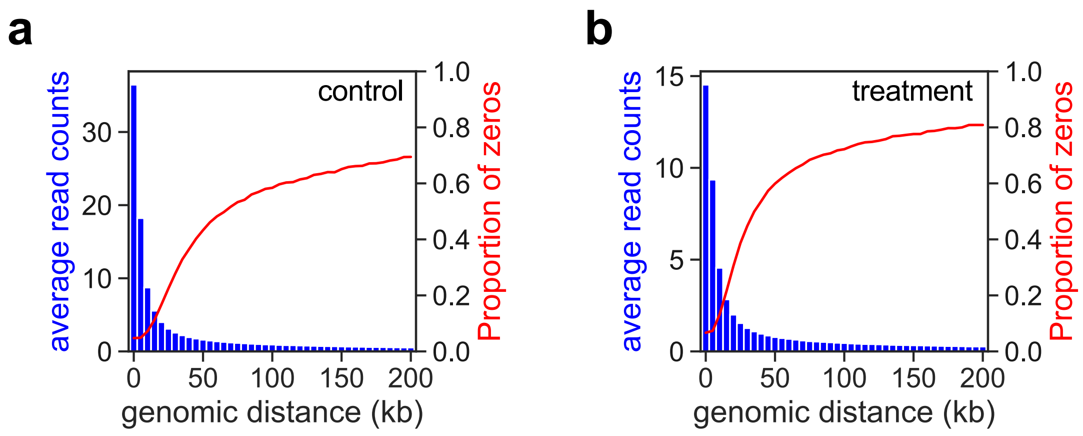


**Supplementary Figure S1. Hi-C read count negatively correlates with genomic distance between bin pairs.** Average read counts and percentage of zeros in all bin pairs at the same genomic distance in two Hi-C datasets. (a) control: GSM2790405; (b) treatment: GSM2790406.

### Supplementary Figure S2

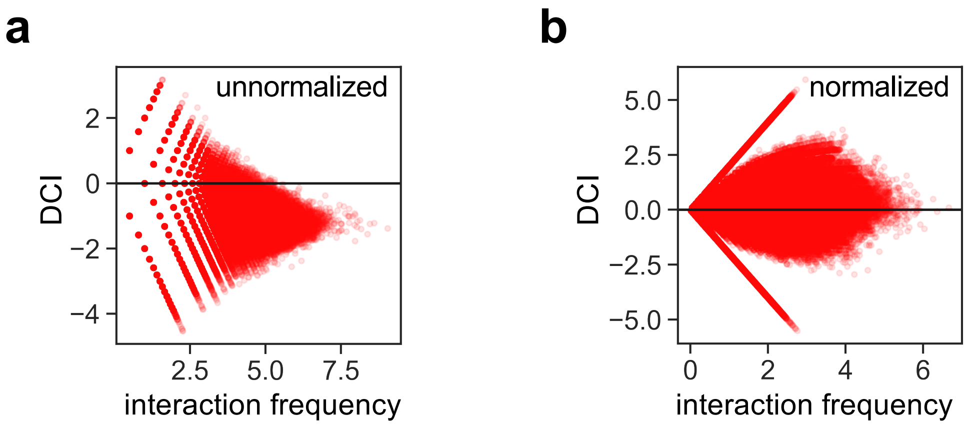


**Supplementary Figure S2. Effect of normalization.** MA plots of averaged interaction frequency (x-axis) and differential chromatin interaction (DCI, y-axis) with unnormalized (a) and normalized (b) Hi-C contact matrices between treatment (GSM2790406) and control (GSM2790405).

### Supplementary Figure S3

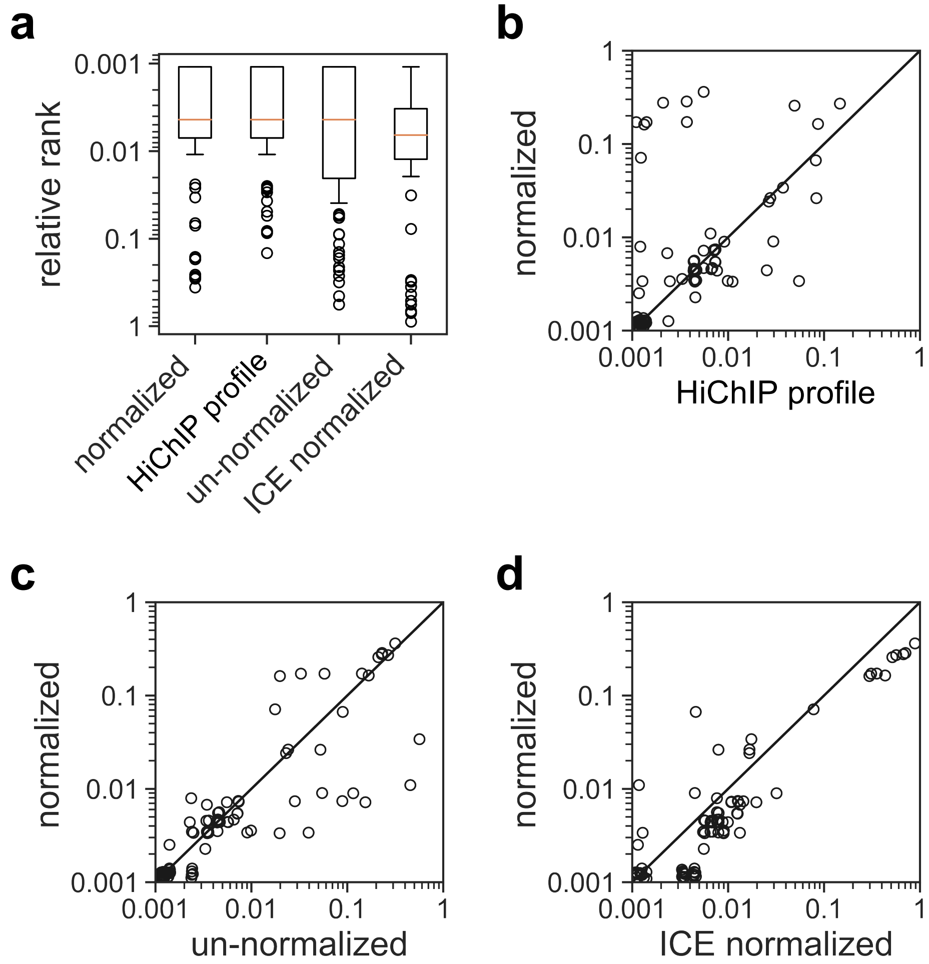


**Supplementary Figure S3. . Comparison of different normalization methods.**

(**a**) BART results of target TRs from 84 HiChIP datasets using different normalization methods. For distance-based normalized (labeled as “normalized”), unnormalized, and ICE normalized, BART was applied to a genomic region profile scored by summarizing the interaction frequencies of each 5kb bin to its flanking bins within 500kb. For the HiChIP profile, BART was applied to the HiChIP sequence read bam file (as positive control). Relative rank represents the rank of the target TR divided by the total number of TRs in the BART library. Center line in the box represents median.

(**b-d**) Comparison of the relative ranks of target TRs in the BART results generated from distance-based normalization against other methods: (**b**) HiChIP profile, (**c**) unnormalized and (**d**) ICE normalization. Each dot represents a dataset from the 84 HiChIP samples. More dots located below the diagonal line indicates that distance-based normalization (y-axis) yields to higher rank in the BART result.

### Supplementary Figure S4

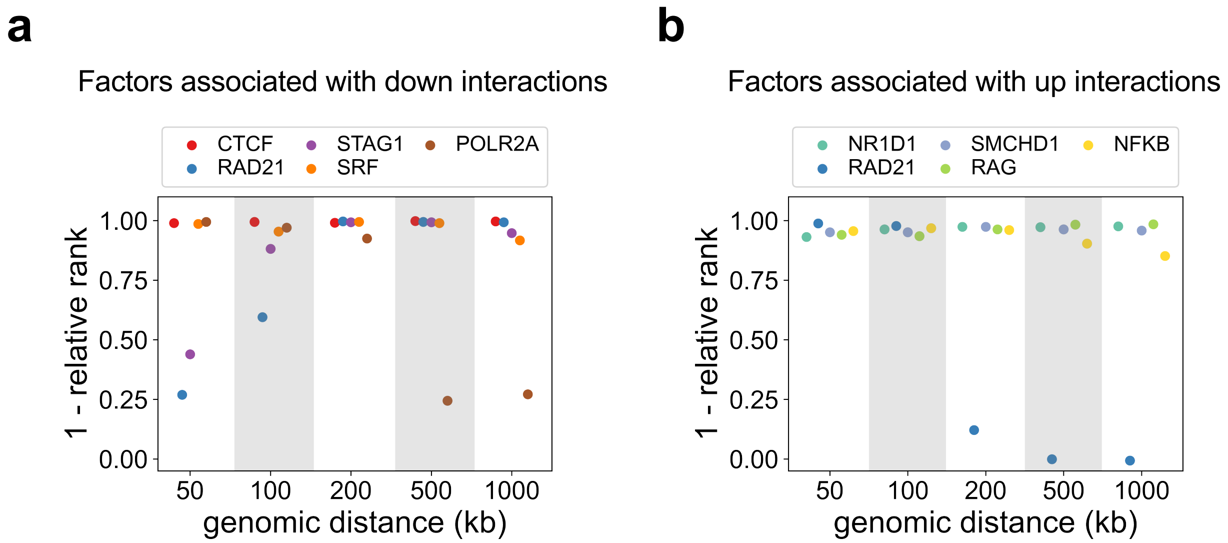


**Supplementary Figure S4. BART3D results on dynamic Hi-C datasets with TR perturbation under different genomic distance settings.** The data were separated for TRs associated with decreased (**a**) and increased (**b**) chromatin interactions. The 1 - relative rank of the perturbed TR in BART3D results were shown for each dataset. Higher scores correspond to higher ranked TRs.
